# Computational approach to identifying universal macrophage biomarker

**DOI:** 10.1101/807347

**Authors:** Dharanidhar Dang, Sahar Taheri, Soumita Das, Pradipta Ghosh, Lawrence S. Prince, Debashis Sahoo

## Abstract

Macrophages are a type of white blood cell, of the immune system, that engulfs and digests cellular debris, cancer cells, and anything else that does not have the type of proteins specific to healthy body cells on its surface. Understanding gene expression dynamics in macrophages are crucial for studying human diseases. Recent advances in high-throughput technologies have enabled the collection of immense amounts of biological data. A reliable marker of macrophage is essential to study their function. Traditional approaches use a number of markers that may have tissue specific expression patterns. To identify universal biomarker of macrophage, we used a previously published computational approach called BECC (Boolean Equivalent Correlated Clusters) that was originally used to identify universal cell cycle genes. We performed BECC analysis on a seed gene CD14, a known macrophage marker. FCER1G and TYROBP were among the top candidates which were validated as strong candidates for universal biomarkers for macrophages in human and mouse tissues. To our knowledge, such a finding is first of its kind.

**CONTRIBUTIONS TO THE FIELD:** We have developed a computational approach to identify universal biomarkers of different entities in a biological system. We applied this approach to study macrophages and identified universal biomarkers of this particular cell type. FCER1G and TYROBP were among the top candidates which were validated as strong candidates for universal biomarkers for macrophages in human and mouse tissues. The expression patterns of TYROBP and FCER1G are found to be more homogeneous compared to currently used biomarkers such as ITGAM, EMR1 (F4/80), and CD68. Further, we demonstrated that this homogeneity extends to all the tissues currently profiled in the public domain in multiple species including human and mouse. FCER1G and TYROBP expression patterns were also found to be extremely specific to macrophages found in various tissues. They are strongly co-expressed together. We believe that these two genes are the most reliable candidates of universal biomarker for macrophages.

## INTRODUCTION

Macrophages are specialized cells involved in the detection, phagocytosis and destruction of bacteria and other harmful organisms. In addition, they can also present antigens to T cells and initiate inflammation by releasing molecules (known as cytokines) that activate other cells. Further, Macrophages migrate to and circulate within almost every tissue, patrolling for pathogens or eliminating dead cells. Critical for immune protection and tissue homeostasis, macrophage functions can be corrupted in multiple disease processes ^1^. Disruption of normal macrophage biology is a hallmark of many diseases, including diabetes^2,3^, asthma^4^, metastatic cancer^5^, tissue fibrosis^6^, and chronic inflammation^6–8^. These characteristics make macrophages a vital element, especially to understand diseases. Further, they are important immune cells that function in tissue repair during homeostasis and in the innate immune response. Inflammation, which can be triggered by infection, is accompanied by a massive expansion of macrophages in affected tissues. The origin of macrophages is thought to be the blood stem cells in the bone marrow. However, a recent study shows that macrophages can initiate cell division and can create a self-replica. These functions are essential to maintain tissue homeostasis^9^. These critical functionalities have propelled researchers to understand macrophages better.

Recent advances in high-throughput sequencing technologies have facilitated large collections of biological datasets. This has propelled significant efforts to model the complexities of macrophage biology. Accordingly, macrophages showed diverse and variable expression patterns, even in the established pool of markers. However, a reliable universal biomarker of macrophages has not been established due to difficulty in experimental techniques and limited purification strategies. Commonly used markers for macrophages such as CD14^10^, ITGAM^11^, CD68^12^ and EMR1^13^ have shown variable expression patterns in different tissues.

Using sequencing data, large scale genomic profiling studies have identified differences in macrophages based on developmental stage, tissue location, and disease process. Novel informatic analysis of these large datasets could leverage the diversity of gene expression data and identify specific patterns and pathways regulating macrophage biology. Collombet et al. have proposed a dynamic logical model of blood cell macrophages using a limited number of gene expression datasets^14^. Such a model may not be generalized as the authors do not consider a wide range of datasets. Boolean modeling has been proposed to study the polarization of macrophages^15,16^. Boolean modeling of the NFκB pathway in bacterial lung infection has been explored.

## MATERIALS AND METHOD

### Data Collection and Annotation

Publicly available microarray databases in Human U133 Plus 2.0 (n=25,955, GSE119087), Mouse 430 2.0 (n=11,758, GSE119085) Affymetrix platform were downloaded from National Center for Biotechnology Information (NCBI) Gene Expression Omnibus (GEO) website ^17–19^. Gene expression summarization was performed by normalizing each Affymetrix platform by RMA (Robust Multichip Average)^20,21^. One hundred ninety-seven published macrophage samples from seven series assayed on the Human U133 Plus 2.0 (GPL570), Human U133A 2.0 (GPL571) and Human U133A (GPL96) platforms were re-analyzed and deposited in GEO with accession no GSE134312. RMA was used to normalize the macrophage CEL files using a modified CDF file that contains the shared probes among the three different platforms. The global human dataset GSE119087 included 106 macrophage samples from GSE134312 dataset. Mouse dataset GSE119085 was also annotated with 327 macrophage samples that were deposited in GEO with accession no GSE135324. In addition to the above training datasets, several human and mouse validation datasets were assembled from GEO. We validate our markers in 39 distinct highly purified mouse hematopoietic stem, progenitor, and differentiated cell populations covering almost the entire hematopoietic system: Gene Expression Commons (GEXC, GSE34723, n = 101)^22^. In addition to GEXC, we also used ImmGen datasets that are also purified mouse blood cells (GSE15907 and GSE127267)^23,24^.

We put together four purified human macrophage datasets: (GSE35449, n=21)^25^, (GSE85333, n=185)^26^, (GSE46903, n=384)^27^, (GSE55536, n=33)^28^.

GSE35449 (PBMC): CD14+ monocytes were isolated from Peripheral blood mononuclear cells (PBMC) using CD14-specific MACS beads and cultured in 6-well plates in media and provided various stimuli: IFN-γ, TNF-α, ultrapure LPS, IL-4, IL-13, or combinations thereof.

GSE85333 (PBMC): Primary human CD14+ monocytes were isolated from the whole blood of 6 donors (3 males, 3 females). These were transformed in macrophages through CSF-1 stimulation over a week. Cells were then subject to an inflammatory stimulus with LPS or IFNa and without any inflammatory stimulus.

GSE46903 (PBMC): Human monocytes were purified from peripheral blood mononuclear cells by MACS, followed by stimulation with GM-CSF or M-CSF for 72 hr.

GSE55536 (iPSDMs and PBMC): Transcriptome analyses of human induced pluripotent stem cell-derived macrophages (IPSDMs) and their isogenic human peripheral blood mononuclear cell-derived macrophage (HMDM) counterparts.

To validate our results in the mouse, we put together four diverse mouse macrophage datasets: (GSE82158, n=163)^29^, (GSE38705, n=511)^30^, (GSE62420, n=56)^31^, and (GSE86397, n=12)^32^.

GSE82158 (interstitial and alveolar): Monocytes, interstitial macrophages, and alveolar macrophages were isolated from naïve mice and RIPK3-/- mice.

GSE38705 (intraperitoneal lavage): Primary macrophages were harvested using four mice per strain which were exposed to either LPS or OxPAPC.

GSE62420 (Brain Microglia): Microglia cells were extracted from 4 regions: cerebellum, cortex, hippocampus, striatum using a magnetic bead-based approach.

GSE86397 (Liver Kupffer cells): Primary Kupffer cells isolated from mouse liver were treated with lipopolysaccharides or IL-4 and the gene expression patterns were analyzed by microarray.

We validated our results on following tissue resident macrophages in human: tumor associated macrophage (GSE117970, n = 116)^33^; lung alveolar macrophages (GSE116560, n = 68)^34^; lung alveolar macrophages (GSE40885, n = 14)^35^; cardiac macrophages (GSE119515, n = 18)^36^; vaginal mucosa and skin macrophages (GSE54480, n = 87)^37^; skin macrophages (GSE74316, n = 77)^38^; peritoneal macrophages (GSE79833, n = 12)^39^; microglia (GSE1432, n= 24)^40^; adipose tissue macrophages (GSE37660, n = 4)^41^.

To validate our results on single cell RNASeq data we use following datasets: mouse inflammatory airway macrophages (GSE120000; n = 1,142)^42^, mouse CX3CR1-derived macrophage from atherosclerotic aorta (GSE123587; n = 5,355)^43^, mouse dissociated whole lung tissue (GSE111664; n = 41,898)^44^, and renal resident macrophages across species (GSE128993; human n = 2,868, mouse n = 3,013, rat n = 3,935, pig n = 4,671)^45^.

We also examined expression patterns in skin Langerhans cell (GSE49475, n = 39)^46^ and dermal dendritic cells (GSE74316, human n = 77, mouse n = 74)^38^.

### StepMiner Analysis

StepMiner is a computational tool that identifies step-wise transitions in a time-series data.^47^ StepMiner performs an adaptive regression scheme to identify the best possible step up or down based on sum-of-square errors. The steps are placed between time points at the sharpest change between low expression and high expression levels, which gives insight into the timing of the gene expression-switching event. To fit a step function, the algorithm evaluates all possible step positions, and for each position, it computes the average of the values on both side of the step for the constant segments. An adaptive regression scheme is used that chooses the step positions that minimize the square error with the fitted data. Finally, a regression test statistic is computed as follows:

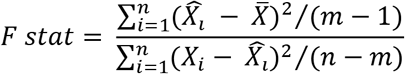

Where *X*_*i*_ for *i* = 1 to *n* are the values, 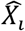 for *i* = 1 to *n* are fitted values. m is the degrees of freedom used for the adaptive regression analysis. 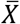 is average of all the values: 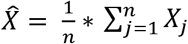. For a step position at k, the fitted values 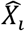 are computed by using 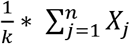 for *i* = 1 to *k* and 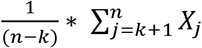 for *i* = *k* + 1 to *n*.

### Boolean Analysis

**Boolean logic** is a simple mathematic relationship of two values, i.e., high/low, 1/0, or positive/negative. The Boolean analysis of gene expression data requires conversion of expression levels into two possible values. The ***StepMiner*** algorithm is reused to perform Boolean analysis of gene expression data.^48^ The **Boolean analysis** is a statistical approach which creates binary logical inferences that explain the relationships between phenomena. Boolean analysis is performed to determine the relationship between the expression levels of pairs of genes. The ***StepMiner*** algorithm is applied to gene expression levels to convert them into Boolean values (high and low). In this algorithm, first the expression values are sorted from low to high and a rising step function is fitted to the series to identify the threshold. Middle of the step is used as the StepMiner threshold. This threshold is used to convert gene expression values into Boolean values. A noise margin of 2-fold change is applied around the threshold to determine intermediate values, and these values are ignored during Boolean analysis. In a scatter plot, there are four possible quadrants based on Boolean values: (low, low), (low, high), (high, low), (high, high). A Boolean implication relationship is observed if any one of the four possible quadrants or two diagonally opposite quadrants are sparsely populated. Based on this rule, there are six different kinds of Boolean implication relationships. Two of them are symmetric: equivalent (corresponding to the highly positively correlated genes), opposite (corresponding to the highly negatively correlated genes). Four of the Boolean relationships are asymmetric and each corresponds to one sparse quadrant: (low => low), (high => low), (low => high), (high => high). BooleanNet statistics (Figure S1A-B) is used to assess the sparsity of a quadrant and the significance of the Boolean implication relationships^48,49^. For each quadrant a statistic S and an error rate p is computed. S > 3 and p < 0.1 are the thresholds used on the BooleanNet statistics to identify Boolean implication relationships.

### BECC (Boolean Equivalent Correlated Clusters) Analysis

BECC analysis is based on Boolean Equivalent relationships, pair-wise correlation and linear regression analysis (Figure S1C). A gene pair was included in the BECC analysis if they had a Boolean Equivalent relationship or both had a Boolean Equivalent relationship with a common third gene. This analysis was performed in two steps. First, a selected probeset of a seed gene was used as a starting point to identify a list of probesets (ProbeSet A) that are Boolean Equivalent to the selected probeset. Next, this list was expanded (ProbeSet B) by identifying other probesets that are Boolean Equivalent to at least one of the probeset from ProbeSet A. Probeset B were further expanded (ProbeSet C, *L*) by repeating the same steps. A score was computed for a pair of probesets from *L* by using the correlation *r* and slope of fitted line *s* (if *s > 1*, *1/s* was used as slope).

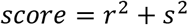

The score is a number between 0 and 2 given r > 0 and s > 0. A matrix of scores *M* was computed for all probesets in *L*. Every row of this matrix was sorted based on the score in ascending order. The whole matrix was then multiplied using a column vector of ranks: *[0 1 2 … len(L)-1]*. In other words, the score for the probeset in row *i gs_i_* was computed as follows:

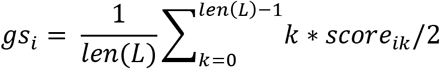

where *score*_*ik*_ is the *k*^*th*^ smallest score for the probeset in row *i*.

StepMiner algorithm was used to compute a threshold to identify the high scoring probesets *gs_i_*. The result of the BECC is this list of high scoring probesets.

### Statistical Justification

Empirical distribution of the pair-wise gene scores were computed for each of our dataset by randomly selecting pairs of probesets. Using this distribution, average probeset score E[gs_i_] and standard deviation can be estimated.

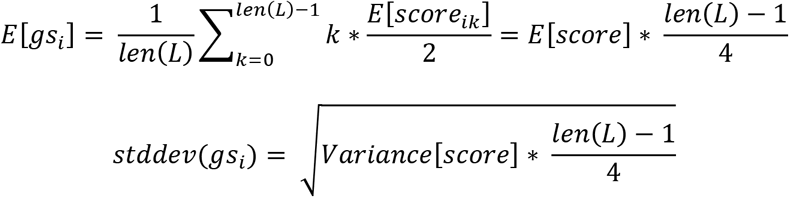

The p-value for the StepMiner identified threshold was computed using a Z-test. All statistical tests were performed using R version 3.2.3 (2015-12-10).

## Results

### BECC identifies macrophage genes in humans

We apply a previously published computational tool called Boolean Equivalent Correlated Clusters (BECC) to mine publicly available gene expression datasets (n = 25,955 human samples, GSE119087)^50^. BECC compares the normalized expression of two genes across all datasets by searching for two sparsely populated, diagonally-opposite quadrants out of four possible quadrants (high-low and low-high), employing the BooleanNet algorithm^48^. The BECC algorithm only focuses on Boolean Equivalent relationships (Figure S1B) to identify potentially functionally-related gene sets (Figure S1C).

To identify potential macrophage genes with this approach, we employed BECC using *CD14* as a seed gene because it is known to be specific for macrophages (Figure 1A)^51,52^. However, CD14 is not considered a universal marker of macrophages because of its variable expression patterns among different types of macrophages^51,52^. Discovering universal biomarkers of a chaotic tissue element such as Macrophage would require suitable datasets of large sizes. We consider publicly available microarray databases in Human U133 Plus 2.0 (n=25,955, GSE119087) Affymetrix platform from GEO.

**Figure 1:**
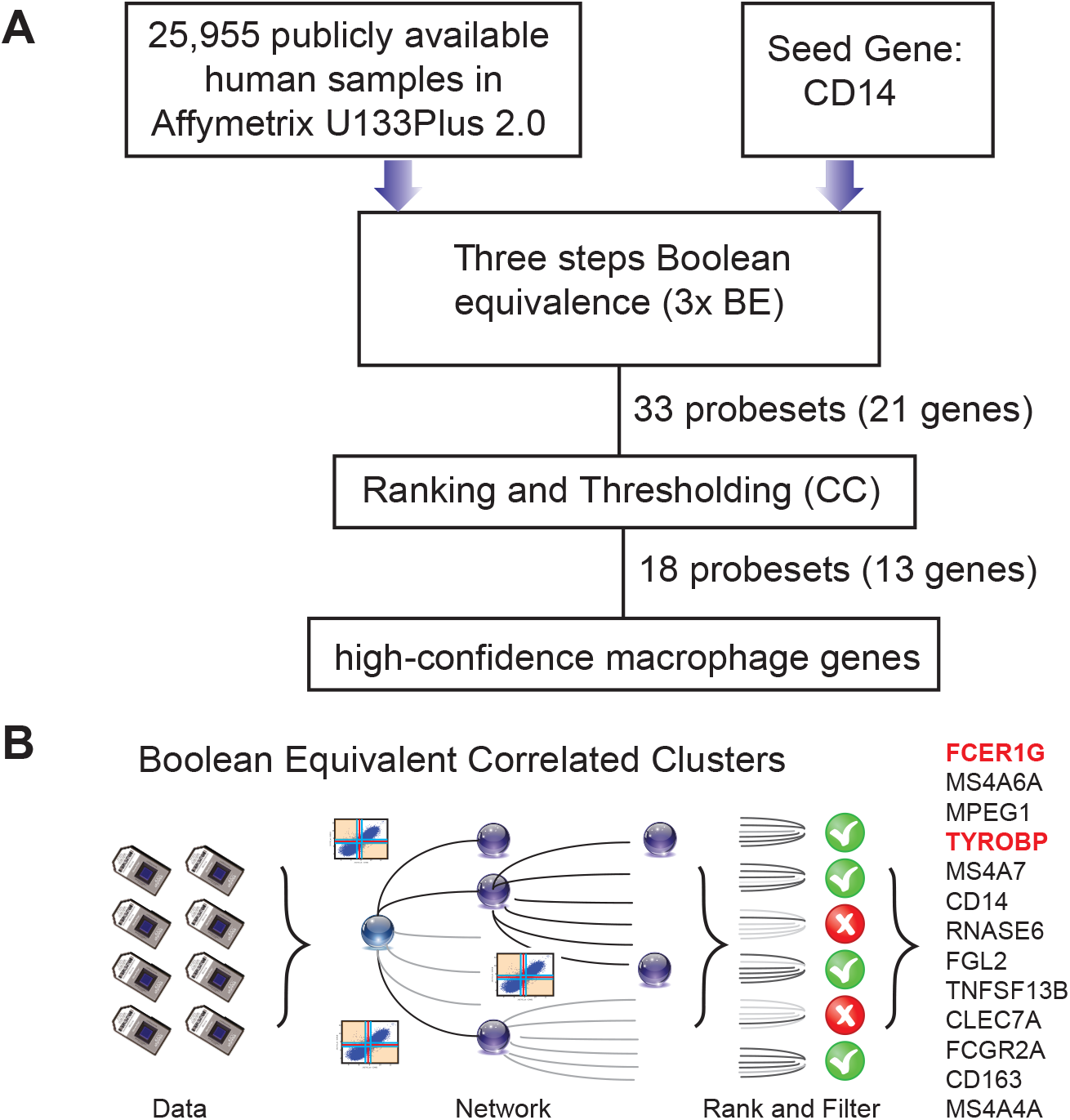
Computational approach to identifying candidate universal macrophage biomarker. (A) A flow chart of the different steps of BECC (Boolean Equivalence Correlated Clusters) to identify robust macrophage biomarker. (B) Overview of BECC illustrating input data, building networks, ranking and filtering that finally selected 13 genes.

The BECC algorithm was first used to identify a set of 9 probesets (ProbeSet A) that were Boolean-Equivalent to the *CD14* gene (201743_at probeset). Then, the same algorithm was used to identify additional probesets that were Boolean-Equivalent to ProbeSet A; pooling the hits in the second step together with those in ProbeSet A resulted in ProbeSet B comprised of 20 probesets. A third step was performed to collect few more candidates resulted in ProbeSet C comprised of 33 probesets (Fig. 1A). BECC computes Boolean Equivalences for three steps because any additional steps have the potential to add significant noise. All probesets in ProbeSet C were then comprehensively analyzed relative to each other to assess the strength of their equivalences. A Boolean-Equivalence score for each probeset within ProbeSet C was computed based on the weighted average of the correlation coefficient and slope in pair-wise analysis with all other probesets, as described in the Methods. This effort resulted in a ranked list of 33 probesets, corresponding to 21 unique genes, based on similarity to *CD14*. The entire ranked list of genes can be accessed online using our web-resource. StepMiner, an algorithm which fits a step function to identify abrupt transitions in series data, was used to compute a threshold on the BE score to identify high-confidence macrophage genes. Imposition of the threshold resulted in the identification of 18 significant probesets, representing 13 unique genes (Fig. 1B). These 13 genes represent candidates for universal macrophage biomarker.

We compared CD14 expression patterns with other known markers such as CD16, CD64, CD68, CD71, CCR5 and ITGAM (Figure S2A-F). CD14 had better dynamic range compared to these other genes. CD71 was weakly correlated with CD14 suggesting that it may have other tissue specific expression patterns. BECC analyses starting with seed genes CD71, and CCR5 returns no results as none of the genes had Boolean equivalent relationships with these genes. CD68 and ITGAM returned too many results, prompted us to increase the threshold (S > 50, p < 0.1) to get specific genes. Finally, we observed that the results from seed gene CD64 had the most overlap with CD14 (Figure S2G). Thus, the BECC results may vary significantly depending on the seed genes. It is better to pick a gene with good dynamic range to get the best answer.

### TYROBP and FCER1G are two strong candidates for universal macrophage biomarker

FCER1G was the top candidate and TYROBP was the fourth candidate based on the BECC-ranking (Fig. 1B). All 13 gene candidates were evaluated on the human and mouse macrophage datasets. FCER1G and TYROBP emerged as a clear winner with strong correlated patterns in both human and mouse dataset (Fig. 2A-B). We expect that the target biomarkers for macrophages would be highly expressed in pure macrophages sample. Fig. 2A and 2B show scatterplot of expression values between TYROBP and FCER1G in both human and mouse respectively with the pure macrophage samples highlighted in red color. We observe that the expression patterns of both TYROBP and FCER1G are high in our carefully annotated macrophage dataset (red color, Fig 2A-B). The orange color samples in Fig. 2A and 2B illustrates rest of the samples from diverse tissue types, including normal, cancer and other diseases. If there are two macrophage-specific genes that are expressed in all macrophage’s subtypes in all tissues, their expression pattern would be tightly correlated in bulk tissue datasets because the gene expression values would be proportional to the amount (or number) of macrophages present in each tissue sample. It is evident that their expression pattern is extremely tightly correlated in all bulk gene expression datasets in both human and mouse. This type of expression patterns suggests that TYROBP and FCER1G are expressed in similar contexts in all tissue. We conclude that TYROBP and FCER1G expression patterns are equivalent to each other. It is a well-known fact that macrophages are present in every tissue and the amount of macrophages vary dramatically between diverse tissue samples. Ideally, a gene that is strongly correlated with the amount of macrophages in a tissue can be considered as a candidate for a universal macrophage biomarker. However, it is hard to assess the exact amount of macrophages in every bulk tissue sample. We observe that TYROBP and FCER1G both are highly expressed in pure macrophage samples (red color, Fig. 2) and they are strongly correlated in every tissue samples in human and mouse. Based on this, we hypothesize that TYROBP and FCER1G - are universally expressed in all macrophages. To validate this claim, we proceed to the next step.

**Figure 2:**
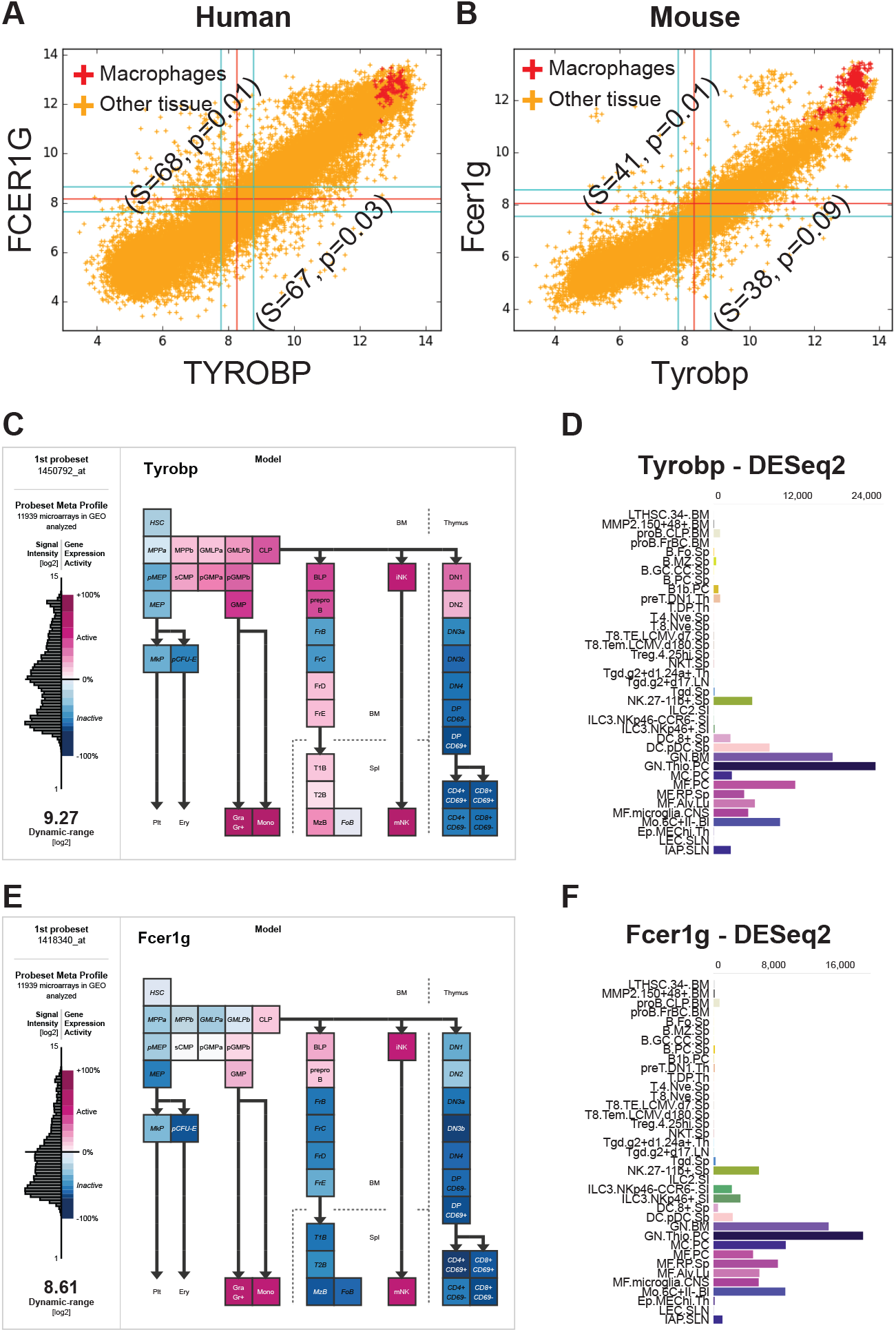
FCER1G and TYROBP expression patterns in human and mouse datasets. (A) A scatterplot of TYROBP and FCER1G in human microarray dataset (n=25,955, GSE119087) with macrophage samples (A subset of GSE134312, n=106) are highlighted in red and the rest of them are in orange color. Every point in the scatterplot is a microarray experiment in Human U133 Plus 2.0 Affymetrix platform. (B) A scatterplot of Tyrobp and Fcer1g in mouse microarray dataset (n=11,758, GSE119085) in Affymetrix Mouse 430 2.0 platform. Similar to panel A, macrophage samples (GSE135324, n=327) are highlighted in red color and the rest of them are in orange color. (C) Expression patterns of Tyrobp in gene expression commons (<u>GEXC</u>). (D) Tyrobp gene expression in Immunological Genome Project (ImmGen) ULI RNASeq dataset (GSE127267) obtained using <u>skyline</u> data viewer from ImmGen website. (E) Expression patterns of Fcer1g in gene expression commons (<u>GEXC</u>). (D) Fcer1g gene expression in ImmGen ULI RNASeq dataset (GSE127267) obtained using <u>skyline</u> data viewer from ImmGen website. (C,E) The data is organized in terms of hematopoietic stem cell differentiation hierarchy and heatmap color code is specified in the figure. (D, F) Gene skyline from ImmGen shows the different purified hematopoietic cell types that were profiled using RNASeq approach.

We have analyzed Tyrobp and Fcer1g expression in GEXC (Fig. 2C, E) and ImmGen ULI RNASeq dataset (Fig. 2D, F). GEXC (Gene Expression Commons) features 39 distinct highly purified mouse blood cells (GSE34723, n = 101)^22^. ImmGen ULI is an open-source project that features expression profiles of the purified immune cell populations^23,24^. We observed that in both of these datasets, the expression patterns of Tyrobp and Fcer1g is exclusively limited to macrophage-like cells and NK cells. This validates our hypothesis that Tyrobp and Fcer1g are universal candidate biomarkers for mouse macrophages.

### FCER1G and TYROBP are highly expressed in purified macrophage datasets

To validate TYROBP and FCER1G as universal biomarkers, we apply pure macrophage datasets collected from several human and mouse tissues (Fig. 3). We put together four purified human macrophage datasets: (GSE35449, n=21)^25^, (GSE85333, n=185)^26^, (GSE46903, n=384)^27^, (GSE55536, n=33)^28^, and four diverse mouse macrophage datasets: (GSE82158, n=163)^29^, (GSE38705, n=511)^30^, (GSE62420, n=56)^31^, and (GSE86397, n=12)^32^.

**Figure 3:**
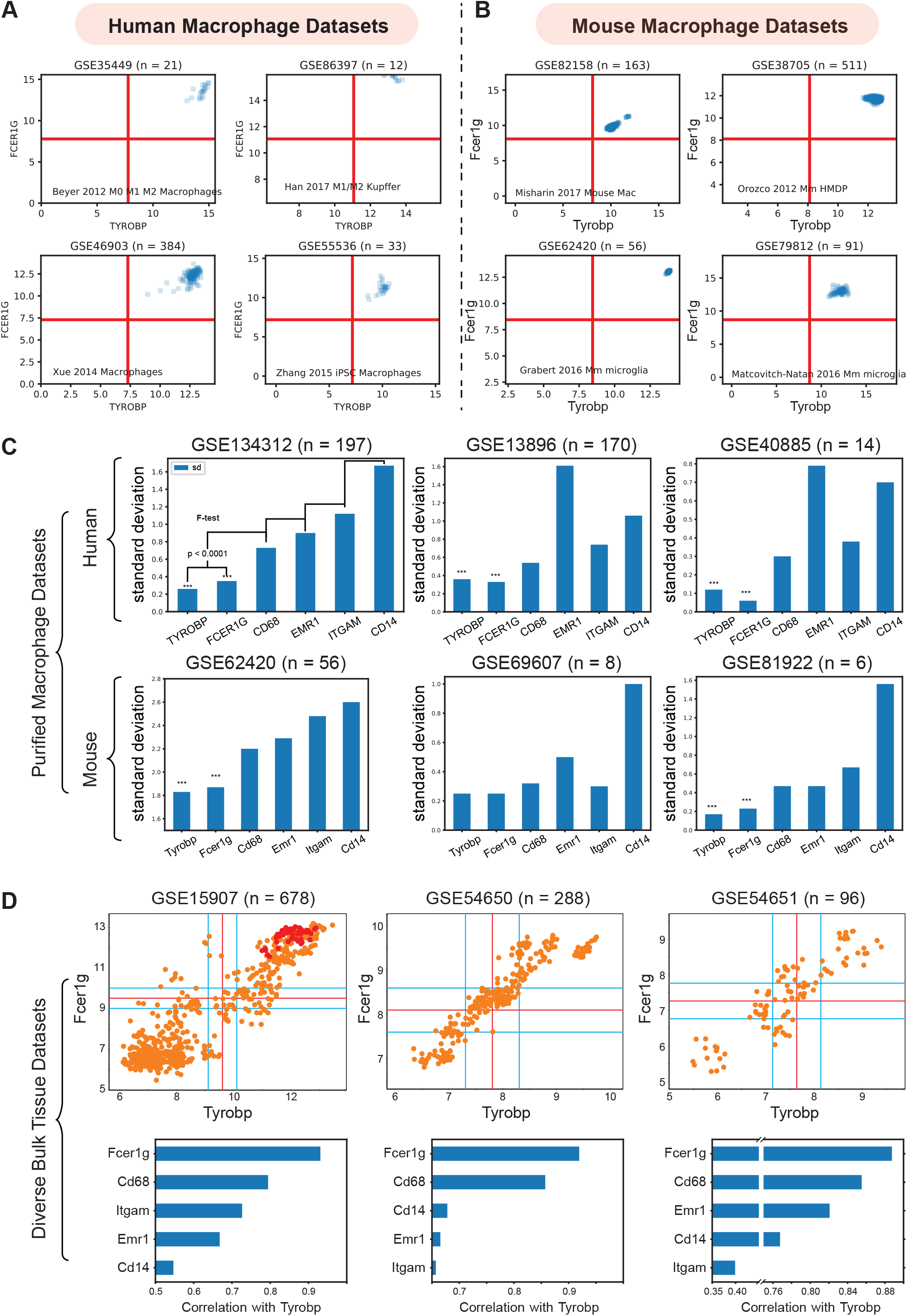
Validation of TYROBP and FCER1G as a universal biomarker of macrophage. (A) Expression patterns of TYROBP and FCER1G in four purified human macrophage datasets: (GSE35449, n=21), (GSE85333, n=185), (GSE46903, n=384), (GSE55536, n=33). (B) Expression patterns of Tyrobp and Fcer1g in four purified mouse macrophage datasets: (GSE82158, n=163), (GSE38705, n=511), (GSE62420, n=56), and (GSE86397, n=12). (C) Standard deviation of TYROBP and FCER1G is compared (F-test) to commonly used macrophage biomarker CD68, EMR1, ITGAM, CD14 in purified macrophage datasets in human and mouse, Only pooled macrophage dataset (GSE134312, n=197) was part of training data and the rest are independent validation dataset. (D) Pearson’s correlation analysis of Fcer1g, Cd68, Emr1, Itgam, Cd14 with Tyrobp shown as a barplot below the scatterplot between Tyrobp and Fcer1g in three independent bulk tissue datasets. Red colored points represent purified macrophage samples while the orange points represent other cell of tissue types.

We analyzed the diverse human and mouse purified macrophage datasets mentioned above. For each microarray or RNASeq dataset, we computed the range of values observed for different genes and assigned the limits of the x and y-axis accordingly. The red lines in each plot represent the middle of the range which are used as a threshold to separate high and low values. As shown in Fig.3A-B, all the samples have high-high expression patterns for both TYROBP and FCER1G. This experiment validates our hypothesis that TYROBP and FCER1G are candidate biomarkers for human and mouse macrophages.

To check if TYROBP and FCER1G are expressed in tissue resident macrophages in human, we analyzed nine other datasets (Figure S3): (A) tumor associated macrophage (GSE117970, n = 13)^33^; (B) lung alveolar macrophages (GSE116560, n = 68)^34^; (C) lung alveolar macrophages (GSE40885, n = 14)^35^; (D) cardiac macrophages (GSE119515, n = 18)^36^; (E) vaginal mucosa and skin macrophages (GSE54480, n = 70)^37^; (F) skin macrophages (GSE74316, n = 12)^38^; (G) peritoneal macrophages (GSE79833, n = 12)^39^; (H) microglia (GSE1432, n= 24)^40^; (I) adipose tissue macrophages (GSE37660, n = 2)^41^. In all cases, we observed have high-high expression patterns for both TYROBP and FCER1G.

We observed differences in expression patterns with respect to skin Langerhans cells (LCs) which are thought to be part of the mononuclear phagocyte system and it is reasonable to classify LCs in the macrophage lineage ^53^. We observed that FCER1G expression is low and TYROBP is high in some human skin LCs (Figure S4A-B): (A) human skin Langerhans cells (GSE49475, n = 9)^46^; (B) human skin Langerhans cells (GSE74316, n = 13)^38^. However, mouse skin LCs showed high-high expression patterns for both Tyrobp and Fcer1g (GSE74316, n = 5)^38^. Dendritic cells (DC) are also mononuclear phagocytes which has lymphoid origin. We also observed that certain human dermal DCs (CD141+) present variable expression patterns with respect to FCER1G (GSE74316, n = 7)^38^. Despite heterogeneity in FCER1G expression patterns, TYROBP expression patterns remain high in most mononuclear phagocytes.

### FCER1G and TYROBP performed better compared to ITGAM, CD68, EMR1

ITGAM^11^, CD68^12^, and EMR1 (F4/80)^13^ are currently established universal biomarkers for macrophages. We analyzed gene expression patterns for the above genes and compared with TYROBP and FCER1G. Our hypothesis is that a universal biomarker should have stable gene expression patterns in pure macrophage samples. We tested this hypothesis in our pooled human macrophage cohorts (GSE134312, n = 197) by measuring the standard deviation of gene expression patterns (Fig. 3C). We observed that TYROBP and FCER1G both have significantly (p < 0.0001) lower standard deviation compared to the established biomarkers. However, since this dataset was part of training data for this analysis, we demonstrated this phenomena in two other independent human datasets GSE13896 (n = 170)^54^, and GSE40885 (n = 14)^35^, and three other mouse datasets GSE62420 (n = 56)^31^, GSE69607 (n = 8)^55^, and GSE81922 (n = 6)^56^. This suggests that macrophages have variable expression patterns for the established biomarkers. However, TYROBP and FCER1G have stable, high, and fairly homogeneous expression patterns in diverse macrophage samples. To further demonstrate the homogeneity, we performed Pearson’s correlation analysis (Fig. 3D) of Tyrobp and Fcer1g in three independent mouse dataset with different tissue and cell types: GSE15907 (microarray, n = 678)^23^, GSE54650 (microarray, n = 288)^57^, GSE54651 (RNASeq, n = 96)^57^. Additionally, a comparison of Fcer1g, Cd68, Emr1, Itgam, and Cd14, revealed that Fcer1g remained the top correlated genes with Tyrobp in these three diverse mouse bulk tissue datasets (Fig. 3D).

### FCER1G and TYROBP is highly expressed in macrophage single cell RNASeq data

We examined expression patterns of FCER1G and TYROBP in several publicly available single cell RNASeq datasets (Figure 4): (A) renal resident macrophages across species (GSE128993; human n = 2,868, mouse n = 3,013, rat n = 3,935, pig n = 4,671)^45^, (B) mouse CX3CR1-derived macrophage from atherosclerotic aorta (GSE123587; n = 5,355)^43^, (C) mouse inflammatory airway macrophages (GSE120000; n = 1,142)^42^, and (D) mouse dissociated whole lung tissue (GSE111664; n = 41,898)^44^. We computed the percentage of single cell sample shows high-high expression patterns with respect to both FCER1G and TYROBP. Renal resident macrophages show 81%, 91%, 97% and 85% in human, mouse, rat, and pig respectively (Figure 4A). Mouse CX3CR1-derived macrophages from atherosclerotic aorta and inflammatory airway macrophages shows 98% (Figure 4B) and 92% (Figure 4C) high-high respectively. However, single cell RNASeq data from dissociated mouse whole lungs show 20% high-high, because this sample contains a mixture of cell types including both the epithelial cells and the macrophages. We computed the percentage of samples that demonstrate high expression pattern for all 13 genes identified by BECC analysis with seed gene CD14, and the common macrophage genes such as CD16 (FCGR3A), CD64 (FCGR1A), CD68, CD71 (TFRC), CCR5, EMR1, ITGAM, in the single cell RNASeq datasets (Figure 4E). We observed that TYROBP and FCER1G expression patterns are consistently high in all datasets, and other genes show significant heterogeneity in their expression patterns.

**Figure 4:**
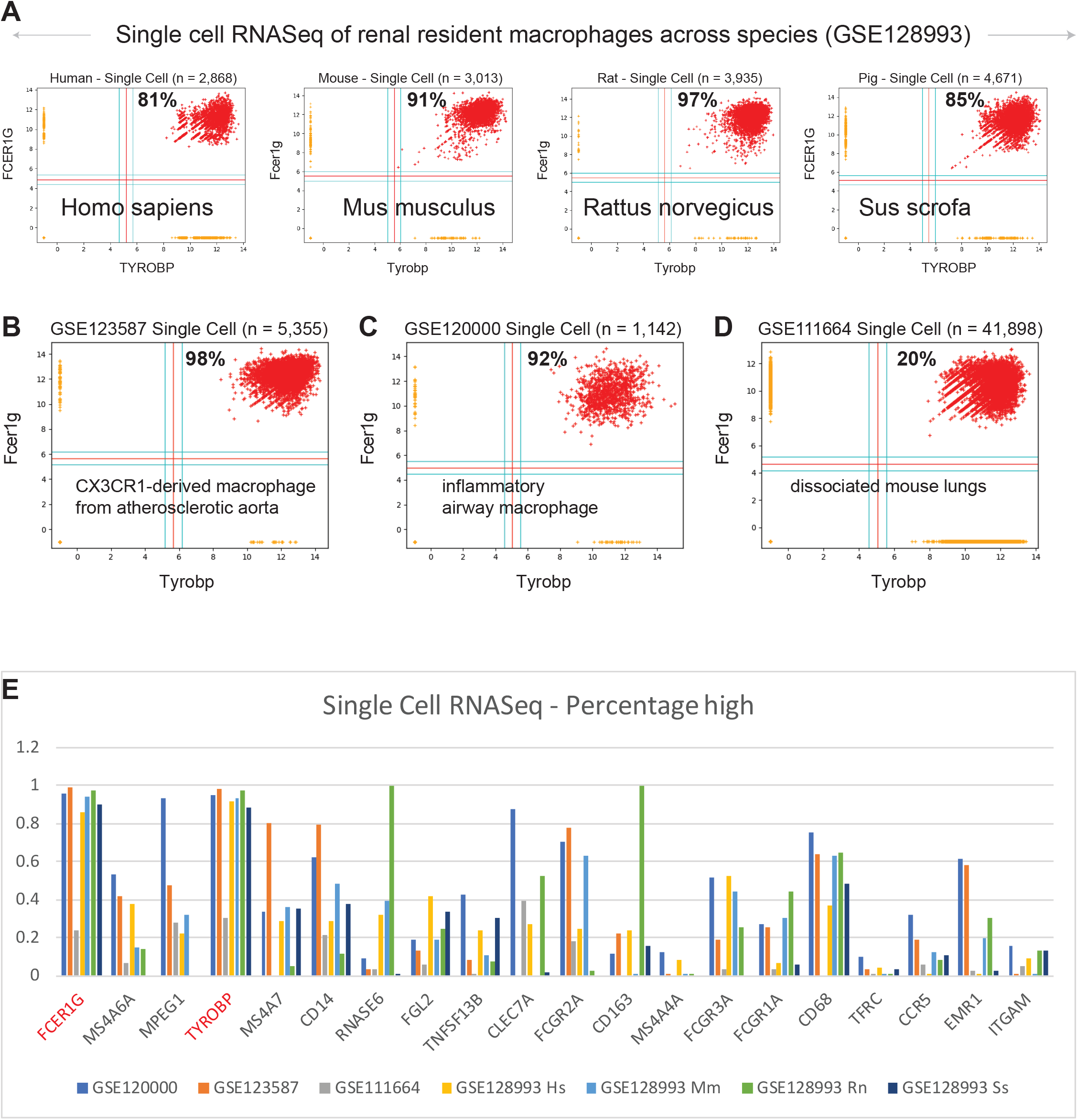
Validation of TYROBP and FCER1G in single cell RNASeq datasets. Scatterplots of expression patterns for TYROBP and FCER1G is shown in several public single cell RNASeq datasets. Red color points denote TYROBP high and FCER1G high samples. Percentage of red points are computed for each scatterplot. Homologous genes are considered for data in mouse, rat and pig. (A) renal resident macrophages across species (GSE128993; human n = 2,868, mouse n = 3,013, rat n = 3,935, pig n = 4,671), (B) mouse CX3CR1-derived macrophage from atherosclerotic aorta (GSE123587; n = 5,355), (C) mouse inflammatory airway macrophages (GSE120000; n = 1,142), and (D) mouse dissociated whole lung tissue (GSE111664; n = 41,898). (E) A bar plot of gene expression values for all 13 genes identified by BECC analysis with seed gene CD14, and the common macrophage genes such as CD16 (FCGR3A), CD64 (FCGR1A), CD68, CD71 (TFRC), CCR5, EMR1, ITGAM, in all the above single cell RNASeq datasets. TYROBP and FCER1G are highlighted in red color.

## DISCUSSION

Normalization is key to perform a reliable high-throughput data analysis. To perform large scale gene expression analysis, all samples from a dataset must be in the same measurement platform. Microarray and RNASeq technologies allow the monitoring of expression levels for thousands of genes simultaneously. However, in these experiments, many undesirable systematic variations are observed even in replicated experiments. Normalization is the process of removing some sources of variation which affect the measured gene expression levels. It is easier to normalize microarray data in one platform. It is much harder to normalize data across platform because it may provide platform-related technical bias. We have pooled all publicly available Affymetrix datasets in U133A, U133A_2 and U133 Plus 2.0 platform for human samples, and in Affymetrix Mouse Genome 430 2.0 Array for mouse samples. We normalized all Affymetrix microarrays using RMA (Robust Multiarray Average) in their respective platforms separately^20,21^. However, Affymetrix datasets in U133A, U133A_2 and U133 Plus 2.0 were pooled into one dataset by using a modified CDF file that contains shared probesets from these three different platforms.

We have developed a computational approach that is geared towards identifying genes that are expressed in macrophages in diverse and almost all context. However, the choice of seed genes can switch gears towards identifying macrophage differentiation and polarization markers such as M1 or M2 phenotypes^58^. Therefore, the results are somewhat sensitive to the choice of seed genes. Seed genes must have good dynamic range and macrophage specificity to perform well. Details of the method, source code and working principles can be found in Figure S1. The method filters out asymmetric relationships (Figure S2A, CD14 vs CD16 is an example) and focus only on the symmetric relationships by using Boolean Implication analysis. In contrast, traditional approach that are purely based on correlation coefficient or linear regression cannot distinguish symmetric vs asymmetric relationships. A macrophage differentiation marker will likely define a subset of macrophages and therefore, in the scatterplot between these genes in Y axis and a universal marker in X axis they may follow asymmetric Boolean Implication: X low => Y low or Y high => X high.

Using CD14 as seed gene, we discovered TYROBP (TYRO protein tyrosine kinase-binding protein) and FCER1G (Fc fragment of IgE receptor Ig) as best candidate for robust universal markers of macrophages. TYROBP is an adapter protein which non-covalently associates with activating receptors found on the surface of a variety of immune cells to mediate signaling and cell activation following ligand binding by the receptors ^59–61^. Interaction of an allergen with FCER1G triggers cell activation, which induces the release of numerous mediators that are responsible for allergic manifestations^62^. Extremely tight correlation is observed between these two genes in all human and mouse microarray datasets (Figure 2A-B). In the GEXC dataset that contain 39 highly purified cell subsets of the mouse blood, Tyrobp and Fcer1g expression were high in the macrophages and the NK cells (Figure 2C, E). B cell and T cell progenitors also show slightly higher expression patterns for Tyrobp and Fcer1g compared to other cell subset such as hematopoietic stem cell (HSC), megakaryocyte (MkP) and erythrocyte (pre-CFU-E) progenitors. Immgen skyline data viewer restricted Tyrobp and Fcer1g expression patterns to granulocytes, microglia and macrophages (Figure 2D, F). Immgen data show low expression in natural killer (NK) and dendritic cells (DCs). Both PBMC-derived and tissue resident macrophages show high expression for TYROBP and FCER1G in diverse settings including single-cell data adding significant strength to our hypothesis (Figure 3 and 4). TYROBP and FCER1G emerge as a winner in direct head-to-head comparison with all 13 genes identified by BECC using CD14 as seed gene, and common macrophage markers such as CD16, CD64, CD68, CD71, CCR5, EMR1 and ITGAM (Figure 4D). One exception was found in human skin Langerhans cells and dermal dendritic cells which show FCER1G low and TYROBP high (Figure S4). This suggest that TYROBP is superior to FCER1G in identifying all mononuclear phagocytes in human irrespective of their lymphoid or myeloid origin. Further validation needed to establish TYROBP and FCER1G as universal marker of macrophages. Literature review showed a computational approach named correlation-based feature subset (CFS) identified TYROBP as part of the hub genes in kidney cancer samples using protein-protein interaction network ^63^. Another study reported that microglia in IDH-mutants are mainly pro-inflammatory, while anti-inflammatory macrophages that upregulate genes such as FCER1G and TYROBP predominate in IDH-wild type GBM ^64^. Tyrobp and Fcer1g was found to be differentially expressed in Alzheimer’s disease (AD) mouse models that demonstrated strong correlation between cortical Aβ amyloidosis and the neuroinflammatory response ^65^. FCER1G was part of a hub gene in a meta-analysis of lung cancer samples ^66^.

Macrophage dysfunction can lead to many human diseases and pathologies, including impaired wound healing, fibrosis^6^, chronic inflammatory diseases^6–8^, diabetic complications^2,3^, and cancer^5^. They play central roles during development^67^, homeostatic tissue processes^1^, tissue repair^1^, and immunity^68^. Macrophages play a vital role in chronic inflammatory diseases such as atherosclerosis^7^ and chronic kidney disease^69^. Due to their large involvement in the pathogenesis of several types of human diseases, macrophages are considered to be relevant therapeutic targets^70^. Macrophage biology, mechanisms of action, and activation phenotypes have been studied extensively in the last few years. Macrophages have a strong tendency to adapt to the microenvironment and to rapidly change in response to environmental stimuli. Thus, it is difficult to design a unique therapeutic strategy based on macrophage modulation that is easily applicable to different kinds of human pathologies. However, our approach appears to identify universal biomarkers that restrict macrophages to a homogeneous state. Our experiments suggest that the variable expression patterns demonstrated by the established macrophage biomarkers is seen both within macrophages and across different tissues. However, in sharp contrast, TYROBP and FCER1G maintain homogeneity of expression patterns in both within macrophages and across different tissues. These candidates would be golden targets of several human diseases as the macrophages would have hard time adapt to any intervention that targets their fundamental properties. The proposed method can be applied in other biological context following the success of macrophage targeting.

## Contributions

Debashis Sahoo – Conceptualization, Data curation, Computation, Formal analysis, Investigation, Methodology, Project administration, Validation, Visualization, Writing – original draft, Writing – review & editing, Funding acquisition, Resources, Supervision

Lawrence S. Prince – Writing – review & editing, Funding acquisition, Resources,

Pradipta Ghosh – Data curation, Analysis, Validation, Writing – review & editing, Funding acquisition, Resources

Soumita Das – Data curation, Validation, Writing – review & editing, Funding acquisition, Resources

Sahar Taheri – Data curation, Validation, Writing.

Dharanidhar Dang – Coordination, Data curation, Investigation, Analysis, Validation, Writing.

## ACKNOWLEGEMENTS

This work was supported by the National Institutes of Health (NIH) grant #R00-CA151673 to DS, 2017 Padres Pedal the Cause / Rady Children’s Hospital Translational PEDIATRIC Cancer Research Award (Padres Pedal the Cause/RADY #PTC2017) to DS, 2017 Padres Pedal the Cause /C3 Collaborative Translational Cancer Research Award (San Diego NCI Cancer Centers Council (C3) #PTC2017) to DS, NHLBI HL126703 to LP and the Gerber Foundation 20180324 to LP.

## Data Submission

All the data generated in the described analyses are submitted to GEO: GSE119085 (mouse), GSE119087 (human), GSE119128 (collections), GSE134312 (human macrophages), GSE135324 (mouse macrophages).

## Data Access

GSE119085 - Mouse Boolean Implication Network GSE119087 - Human Boolean Implication Network

GSE119128 - An unbiased Boolean analysis of public gene expression data for cell cycle gene classification

GSE134312 - Pooled macrophage datasets from GEO GSE135324 - Pooled mouse macrophage datasets from GEO

## ABBREVIATIONS

BECC: Boolean Equivalent Correlated Clusters
GEO: Gene Expression Omnibus
ImmGen: Immunological Genome Project
NCI: National Cancer Institute
NIH: National Institute of Health

---

**Figure S1:**
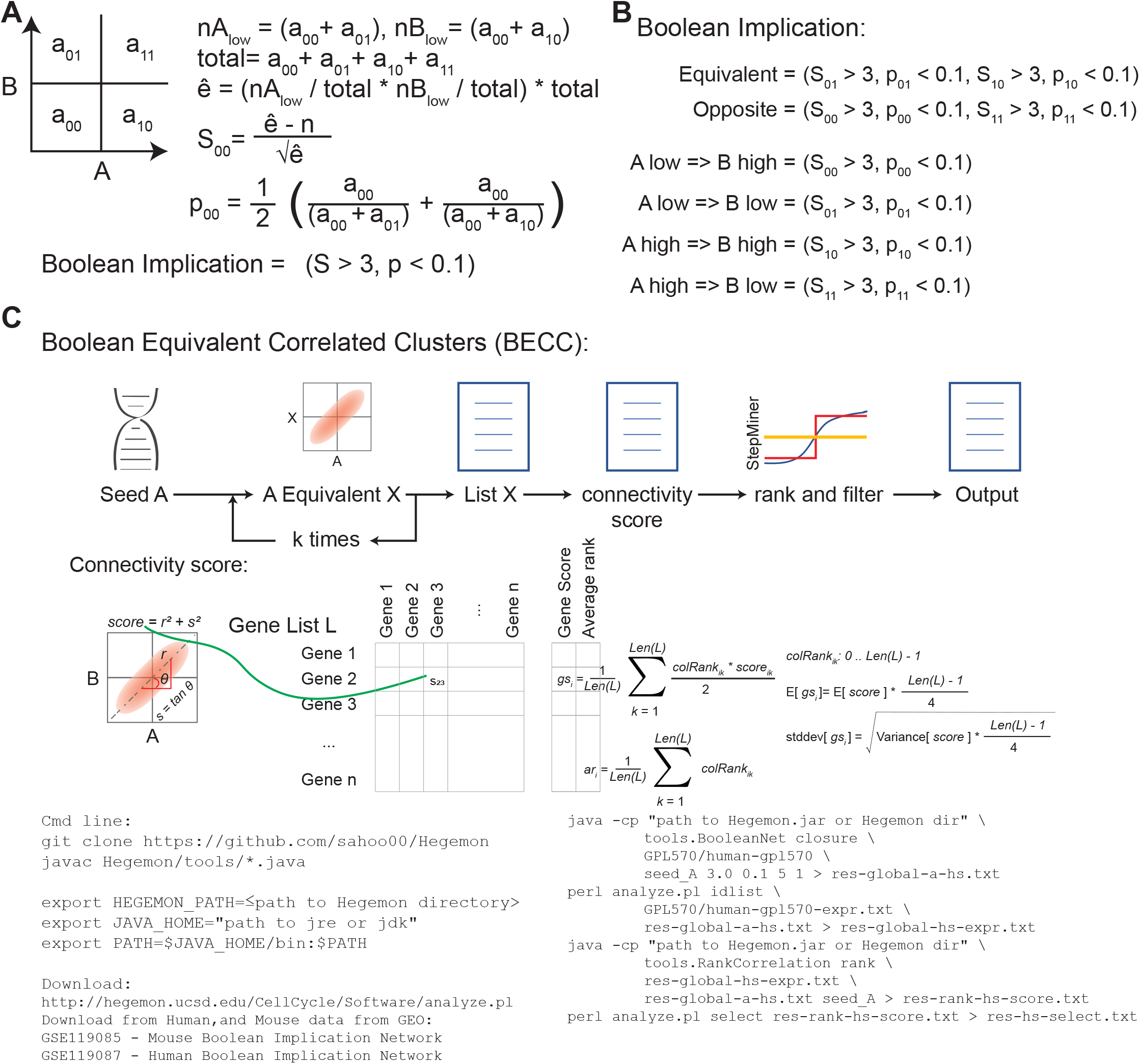
Computational approaches for Boolean analysis. (A) BooleanNet statistic. (B) Deriving Boolean implication relationships using BooleanNet statistic. (C) Workflow and detailed steps of the BECC (Boolean Equivalent Correlated Clusters) tool. A seed gene A is used to extract a list of genes L that are connected by Boolean equivalent relationship directly or indirectly depending on the number k times the loop is considered. A connectivity score is computed for each gene in list L by using the matrix of scores between all pairs that determines how tightly a gene is related to the cluster of genes in L. A gene score is computed as weighted average of the column ranks for each gene. Average gene rank is also computed for each gene which is used to rank genes. StepMiner is used to put a threshold on the gene score to filter highly ranked genes. Output is the candidate gene list computed by BECC.

**Figure S2:**
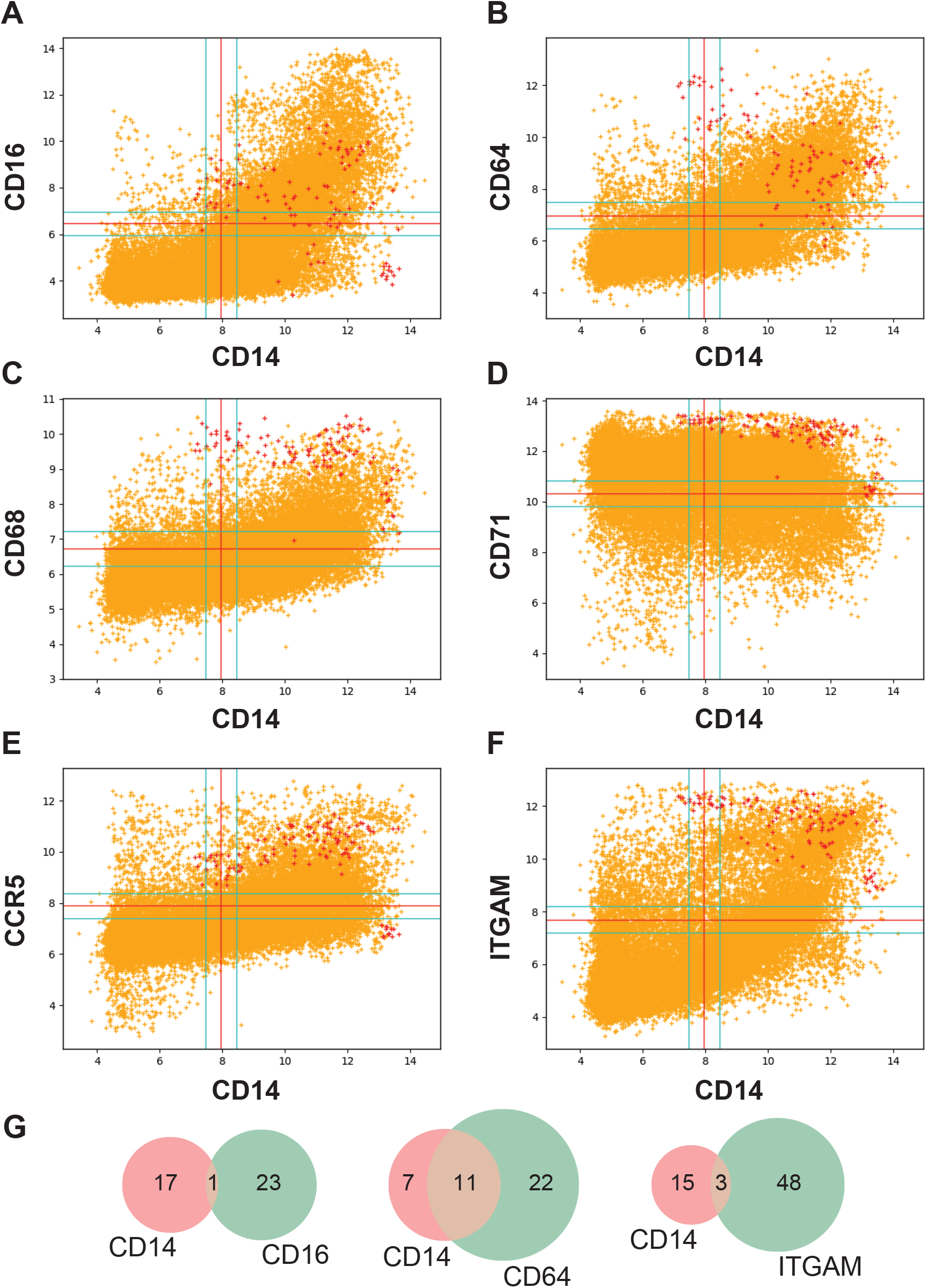
Relationship between CD14 and other known macrophage markers. Scatter plots of gene expression data between CD14 (201743_at) and other known macrophage markers in global human dataset (GSE119087). Red points corresponds to purified macrophage samples (GSE134312). Orange points corresponds to other human samples. (A) CD14 vs CD16 (204006_s_at). (B) CD14 vs CD64 (216950_s_at). (C) CD14 vs CD68 (203507_at). (D) CD14 vs CD71 (208691_at). (E) CD14 vs CCR5 (206991_s_at). (F) CD14 vs ITGAM (205786_s_at). (G) Overlap between the BECC analyses based on different seed genes. BECC analyses on CD71 and CCR5 returned no results. BECC on ITGAM and CD68 returned too many results, therefore we increased BooleanNet statistic to S > 50, p < 0.1 for these two genes.

**Figure S3:**
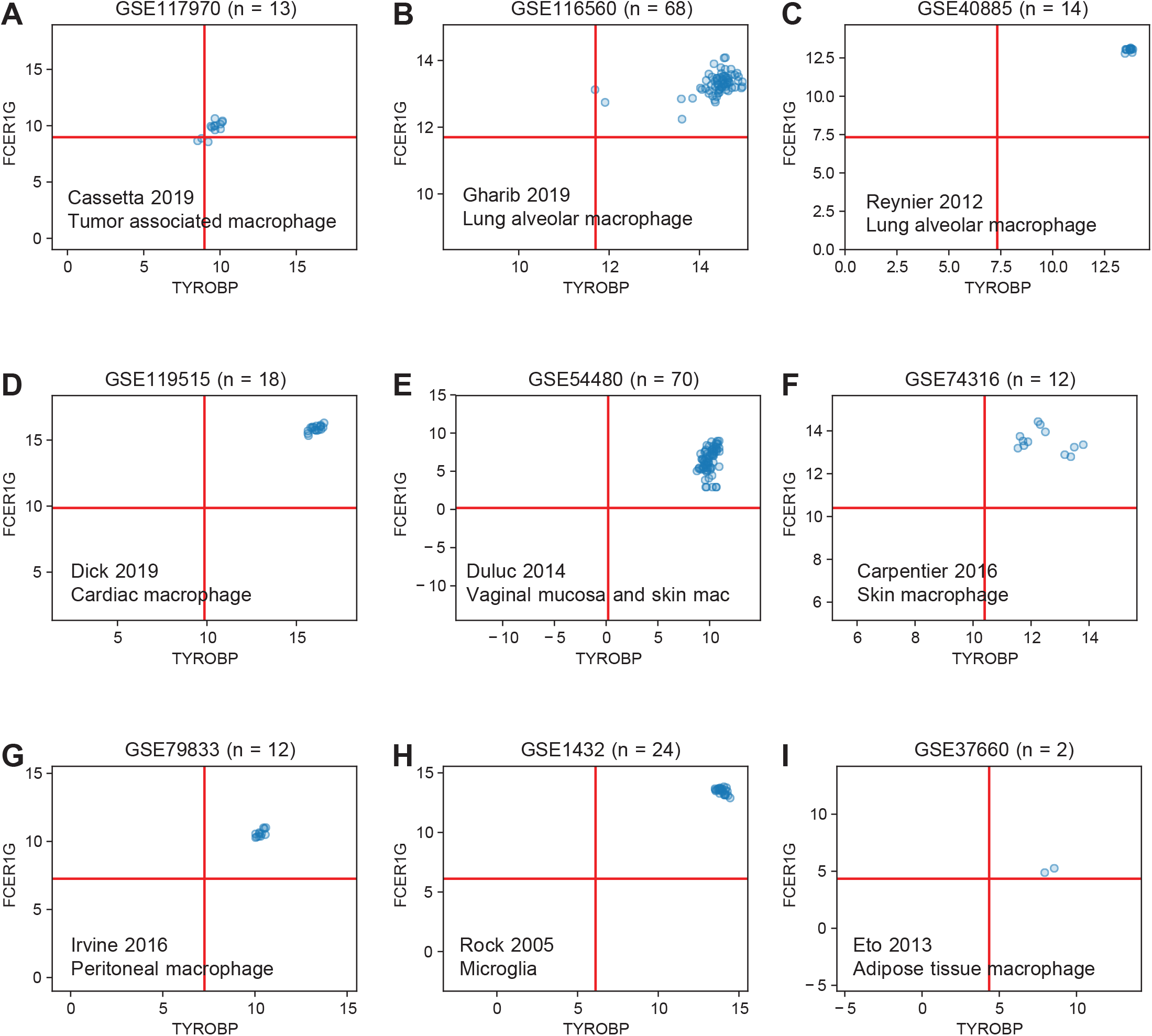
TYROBP and FCER1G expression in human tissue resident macrophages. The limits of the axes were set to the minimum and maximum expression values in each dataset. The red lines denote the mid-point between the minimum and maximum values. Scatter plots of TYROBP and FCER1G in human tissue resident macrophages in nine different context: (A) tumor associated macrophage (GSE117970, n = 13); (B) lung alveolar macrophages (GSE116560, n = 68); (C) lung alveolar macrophages (GSE40885, n = 14); (D) cardiac macrophages (GSE119515, n = 18); (E) vaginal mucosa and skin macrophages (GSE54480, n = 70); (F) skin macrophages (GSE74316, n = 12); (G) peritoneal macrophages (GSE79833, n = 12); (H) microglia (GSE1432, n = 24); (I) adipose tissue macrophages (GSE37660, n = 2).

**Figure S4:**
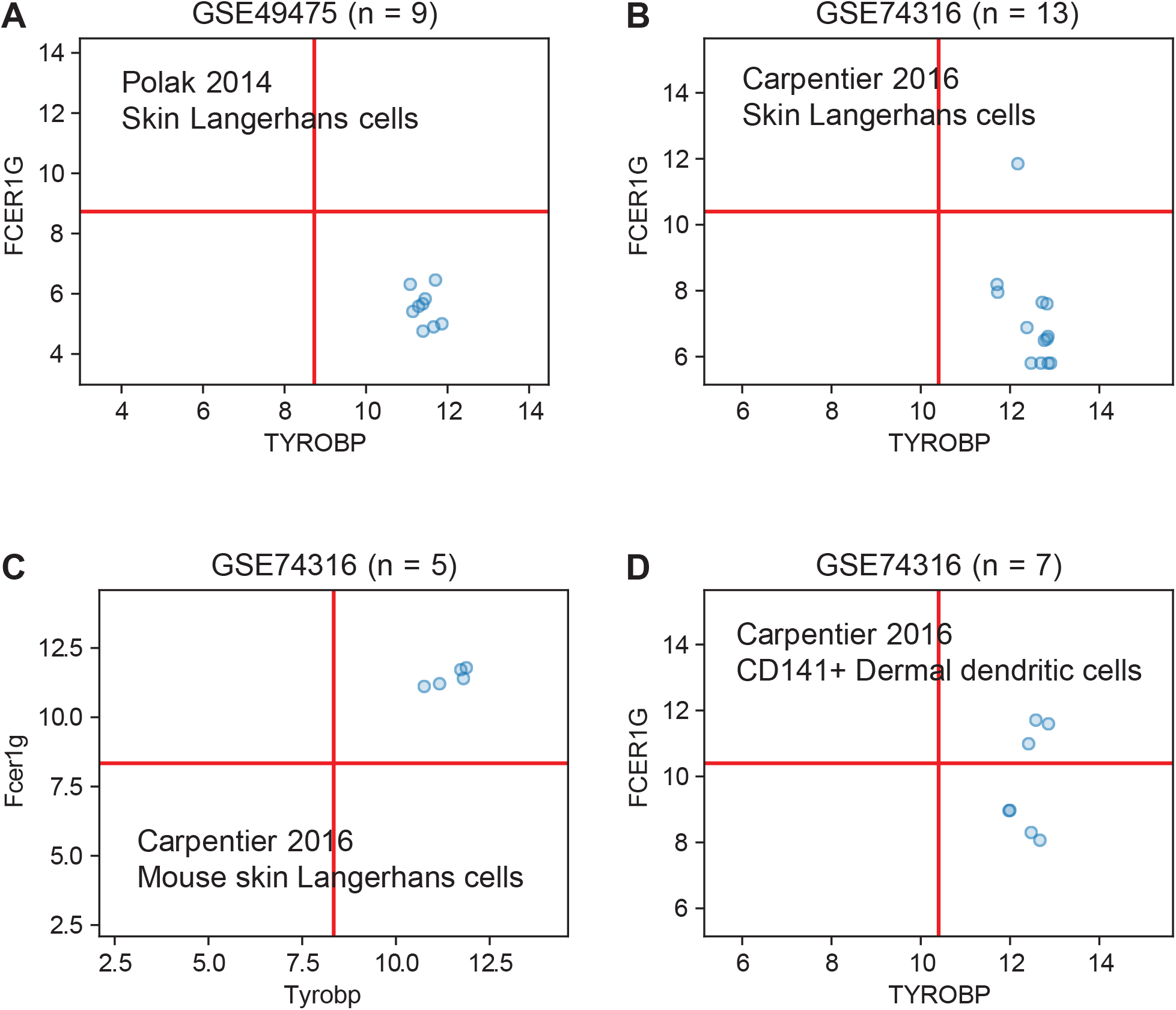
TYROBP and FCER1G expression in skin LCs and DCs. The limits of the axes were set to the minimum and maximum expression values in each dataset. The red lines denotes the mid point between the minimum and maximum values. Scatter plots of TYROBP and FCER1G in skin Langerhans cells and dendritic cells: (A) human skin Langerhans cells (GSE49475, n = 9); (B) human skin Langerhans cells (GSE74316, n = 13); (C) mouse skin Langerhans cells (GSE74316, n = 5); (D) human CD141+ dermal dendritic cells (GSE74316, n = 7).

